# A Behavioral Association Between Prediction Errors and Risk-Seeking: Theory and Evidence

**DOI:** 10.1101/2020.04.29.067751

**Authors:** Moritz Moeller, Jan Grohn, Sanjay Manohar, Rafal Bogacz

## Abstract

Reward prediction errors (RPEs) and risk preferences have two things in common: both can shape decision making behavior, and both are commonly associated with dopamine. RPEs drive value learning and are thought to be represented in the phasic release of striatal dopamine. Risk preferences bias choices towards or away from uncertainty; they can be manipulated with drugs that target the dopaminergic system. The common neural substrate suggests that RPEs and risk preferences might be linked on the level of behavior as well, but this has never been tested. Here, we aim to close this gap. First, we apply a recent theory of learning in the basal ganglia to predict how exactly RPEs might influence risk preferences. We then test our behavioral predictions using a novel bandit task in which value and risk vary independently across options. Critically, conditions are included where options vary in risk but are matched for value. We find that subjects become more risk seeking if choices are preceded by positive RPEs, and more risk averse if choices are preceded by negative RPEs. These findings cannot be explained by other known effects, such as nonlinear utility curves or dynamic learning rates. Finally, we show that RPE-induced risk-seeking is indexed by pupil dilation: participants with stronger pupillary correlates of RPE also show more pronounced behavioral effects.

**Author’s summary:** Many of our decisions are based on expectations. Sometimes, however, surprises happen: outcomes are not as expected. Such discrepancies between expectations and actual outcomes are called prediction errors. Our brain recognises and uses such prediction errors to modify our expectations and make them more realistic--a process known as reinforcement learning. In particular, neurons that release the neurotransmitter dopamine show activity patterns that strongly resemble prediction errors. Interestingly, the same neurotransmitter is also known to regulate risk preferences: dopamine levels control our willingness to take risks. We theorised that, since learning signals cause dopamine release, they might change risk preferences as well. In this study, we test this hypothesis. We find that participants are more likely to make a risky choice just after they experienced an outcome that was better than expected, which is precisely what out theory predicts. This suggests that dopamine signalling can be ambiguous--a learning signal can be mistaken for an impulse to take a risk.

## Introduction

Reward-guided learning in humans and animals can often be modelled simply as reducing the difference between the obtained and the expected reward—a reward prediction error. This well-established behavioral phenomenon (Rescorla and Wagner 1972) has been linked to the neurotransmitter dopamine (Schultz, Dayan et al. 1997). Dopamine neurons project to brain areas relevant for reward learning, such as that striatum, the cortex and the amygdala (Wise 2004, Björklund and Dunnett 2007). Dopamine activity is known to change synaptic efficacy in the striatum (Reynolds, Hyland et al. 2001) and has been causally linked to learning (Steinberg, Keiflin et al. 2013). This and other biological evidence have led to a family of mechanistic theories of learning within the basal ganglia network (Collins and Frank 2014, Mikhael and Bogacz 2016, Möller and Bogacz 2019). According to these models, positive and negative outcomes of actions are encoded separately in the direct and indirect pathways of the basal ganglia.

Crucially, the balance between those pathways is also controlled by dopamine (Gerfen and Surmeier 2011): An increased dopamine level promotes the direct pathway, whereas low levels of dopamine promote the indirect pathway. The family of basal ganglia models includes these modulatory mechanisms too; this makes them consistent with well-studied phenomena whereby dopamine modulates how uncertainty and risk affect decision making: dopaminergic medication can bias human decision making towards or away from risk (Voon, Hassan et al. 2006, Gallagher, O’Sullivan et al. 2007, St Onge and Floresco 2009, Weintraub, Koester et al. 2010), and phasic responses in dopaminergic brain areas predict people’s moment-to-moment risk-preferences (Chew, Hauser et al. 2019).

In summary, ample evidence suggests that dopamine bursts are related to distinct behavioral phenomena—learning and risk-taking—by way of 1) acting as reward prediction errors, affecting synaptic weights during reinforcement learning, and 2) inducing risk-seeking behavior directly. There is no obvious a priori reason for those functions to be bundled together; in fact, one would perhaps expect them to work independently, and their conflation might lead to interactions, unless some separation mechanism exists. There have been different suggestions for such separation mechanisms: it has been proposed that the tonic level of dopamine might modulate behavior directly, while phasic dopamine bursts provide the prediction errors necessary for reward learning (Niv 2007). Alternatively, cholinergic interneurons might flag dopamine activity that is to be interpreted as prediction errors by striatal neurons (Berke 2018). However, it has also been suggested that the prediction errors encoded by dopaminergic neurons might drive both learning and decision making simultaneously (Bogacz 2020).

Curiously, even though the multi-functionality of dopamine has been noted and separation mechanisms have been proposed, interference between the different functions has, to our knowledge, never been tested experimentally. Here, we investigate this: if dopamine indeed provides prediction errors and modulates risk preferences at the same time, do these two processes interfere with each other, or are they cleanly separated by some mechanism? In other words, we test whether prediction errors are associated with risk-seeking.

A common method to provoke prediction-error related dopamine bursts in humans is to present cues and outcomes in sequential decision-making tasks, hence causing prediction errors both when options are presented, and at the time of outcome (Seymour, O’Doherty et al. 2004, Pessiglione, Seymour et al. 2006, Niv, Edlund et al. 2012). To test whether such prediction errors induce risk seeking, we used a learning task in which prediction errors are followed by choices between options with different levels of risk. If there was a clear separation of roles, then risk preferences should be independent of prediction errors.

Incomplete separation, in contrast, should result in a correlation between risk preferences and preceding prediction errors. In particular, we hypothesized that positive prediction errors, occurring when expectations are exceeded, should induce risk seeking, while negative prediction errors should lead to risk aversion.

We further hypothesized that participants who experience stronger prediction errors also show stronger risk preferences. Thus, we tracked participants’ pupil dilation, which is known to reflect surprising events such as prediction errors (Preuschoff, t Hart et al. 2011, Cavanagh, Wiecki et al. 2014, Browning, Behrens et al. 2015, Lawson, Mathys et al. 2017). We expected that increased perception of prediction errors should lead to increased pupil responses as well as stronger risk preferences, and hence predicted a correlation between pupil response to prediction errors and risk seeking associated with prediction errors. Overall, we found effects that were consistent with our predictions: Risk seeking was higher when choices followed positive prediction errors than when they followed negative prediction errors. These preferences emerged gradually over the course of learning and could not be explained by any of several other known mechanisms. In addition, they were associated with pupil dilation in a way predicted by our theory.

## Results

### Task & Theory

In this first section, we introduce our task and provide a detailed theoretical analysis of the behavior we expect from our participants. This analysis is based on models of the basal ganglia network and allows us to derive concrete predictions and models for behavior which we use for data analysis below.

#### Task

Our task consisted of sequences of two-alternative forced choice trials. On each trial, after an inter-trial interval (ITI) of 1 s, two stimuli (fractal images, Fig 1A) were drawn from a set of four stimuli and shown to the participant, who had to choose one. Following the choice, after a short delay of 0.8 s a numerical reward between 1 and 99 was displayed under the chosen stimulus for 1.5 s. Then, the next trial began. Participants were instructed to try to maximize the total number of reward points throughout the experiment.

**Fig 1:**
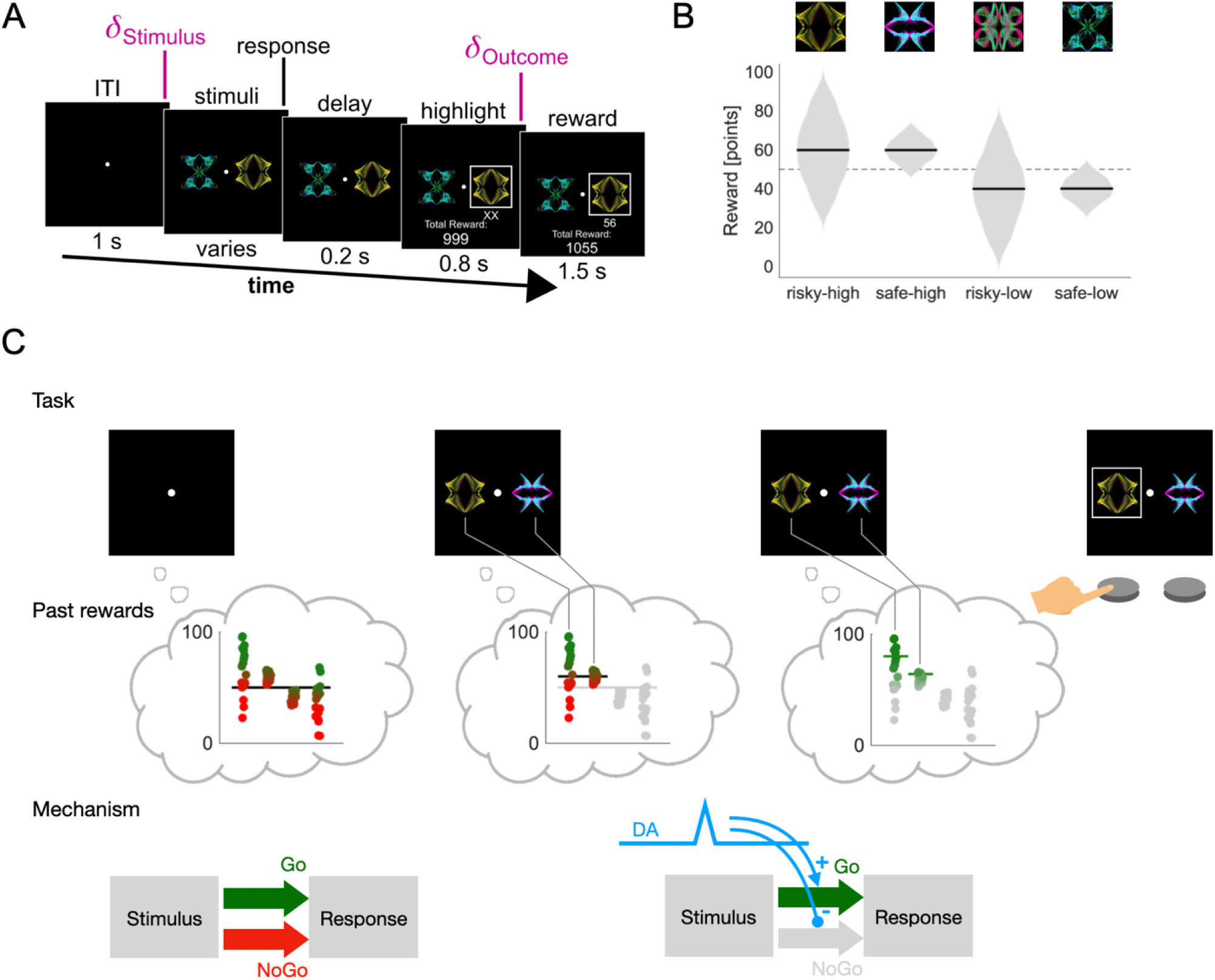
Task design and theory. A) Task structure. Each trial begins with a blank screen. After 1 s, two out of four possible stimuli are selected and shown to the participant. The participant chooses one of them, which is then highlighted after a delay of 0.2 s. After another delay of 0.8 s, a point reward is displayed underneath the chosen stimulus. During each trial two separate prediction errors occur: At stimulus presentation the participant discovers whether the two displayed stimuli are better (positive prediction error) or worse (negative prediction error) than the expected average stimulus. At outcome presentation the participant discovers whether the obtained reward is higher or lower than the expected average reward of the selected stimulus. B) Reward distributions. Each stimulus (top) is associated with a different reward distribution (bottom). The distributions differ in mean (60 points versus 40 points) and standard deviation (20 points versus 5 points). Reward distributions were unknown to participants and the association of stimuli and distributions was shuffled between blocks. The dashed line in the background indicates the middle of the reward range, which was at 50 points. C) Events during the first half of a trial (ITI to response) according to our theory. As the blank screen appears, the participant’s reward expectation (black horizontal line in the first thought-bubble) is based on all past rewards (distributions in the first thought-bubble). Above-average past rewards are encoded in the Go pathway (green dots) and below-average past rewards in the NoGo pathway (red dots). They are weighed equally since the Go and the NoGo pathways are in balance (first diagram in bottom row). As the stimuli appear, only the relevant past rewards are considered (irrelevant rewards greyed out in the middle thought-bubble), and the reward expectation changes (black horizontal line in the middle thought- bubble is higher than the black line in the first thought-bubble). The upward change in reward expectation constitutes a temporal difference prediction error which is signaled by increased dopamine transmission in the striatum (blue transient in bottom row; delta stimulus in panel A). This suppresses the NoGo pathway and enhances the Go pathway (second diagram in bottom row). As a result, below-average past rewards are ignored, and the focus is on above average past rewards (below average rewards greyed out in last thought-bubble). The stimulus with the larger spread is now valued higher (green lines in last thought-bubble) and is therefore chosen.

The reward on each trial depended on the participant’s choice: each stimulus was associated with a specific reward distribution from which rewards were sampled. The four reward distributions associated with the four stimuli were approximately Gaussian and followed a two-by-two design: the mean of the Gaussian could be either high or low (60 or 40), and the standard deviation could be either large or small (20 or 5), resulting in four reward distributions in total (risky-high, risky-low, safe-high and safe-low, Fig 1B). The names derive from the idea that it is “risky” to pick a stimulus associated with a broad reward distribution, since outcomes might deviate a lot from the expected outcome. Correspondingly, it is “safe” to pick a stimulus with a narrow distribution, since the outcomes will mostly be as expected.

We organize trials into three conditions: 1) “**different**”: trials in which the shown stimuli have different average rewards (for example risky-high and safe-low, which have average rewards of 60 and 40 respectively), 2) “**both-high**”: trials in which both stimuli have a high average reward (risky-high and safe-high, both have an average reward of 60 points) and 3) “**both-low**”: trials in which both stimuli have a low average reward (risky-low and safe-low, both have an average reward of 40 points).

#### Theoretical analysis of learning and decision making

In this section, we sketch a mechanistic theory of learning and decision-making in our task. This theory is used to derive the computational model we use in our modelling analysis (see Results/Modelling). We also use it to derive behavioral predictions (see Results/Task & Theory/Behavioral Predictions).

Our theory is based on (Mikhael and Bogacz 2016, Möller and Bogacz 2019). Its premise is that choices are governed by competing action channels in the basal ganglia (BG) network, an assumption common to many models of the basal ganglia (Gurney, Prescott et al. 2001, Frank 2006). We assume that for each option *i* in our task, there is one such action channel, and that the probability of choosing option *i* depends on the total activation *A*_*i*_ of that action channel. There are two contributions to this activation: excitation of magnitude *G*_*i*_ through the direct pathway (also called Go pathway), and inhibition of magnitude *N*_*i*_ through the indirect pathway (also called No-Go pathway). These contributions are differentially modulated by dopamine (DA), mediated through the D1 and D2 receptors expressed in the direct and indirect pathway respectively. Let *δ* represent the deviation of DA levels from baseline (i.e., *δ* = 0 means baseline DA levels, *δ* > 0 means DA is above baseline etc.). Then, the activation of an action channel corresponding to option *i* can be approximated as

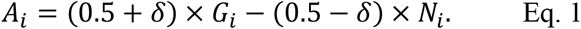

What determines *G*_*i*_ and *N*_*i*_ for each option? According to (Mikhael and Bogacz 2016, Möller and Bogacz 2019), the direct and indirect pathway are subject to DA-dependent plasticity. Hence, *G* and *N* change through reward-driven learning—roughly, *G* tracks the upper end of the reward distribution, while *N* tracks the lower end. More accurately, we may assume that *G* − *N* tracks *Q*, the mean of the reward distribution, according to the rule

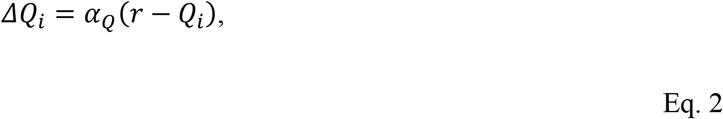

and that *G* + *N* tracks *S*, the spread of the reward, according to the rule

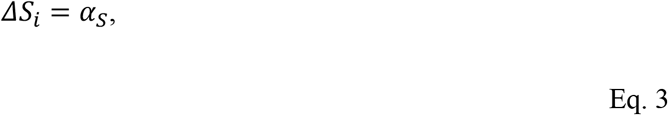

where *α*_*Q*_ > *α*_*S*_. Using this notation, Eq. 1 reads *A*_*i*_ = *Q*_*i*_ + *δ* × *S*_*i*_ — the impact of reward spread on action activation is *gated* by the level of DA.

In our task, the *Q* and *S* of the four action channels should converge to the means and spreads of the four reward distributions. With DA at baseline (i.e., *δ* = 0) activation is proportional to *Q*, and hence to the mean reward. Choices should thus be biased towards options with high mean rewards. If DA levels are increased (i.e., *δ* > 0), the learned spread *S* contributes positively to action activation, biasing choices towards risky options. If, on the other hand, DA levels are below baseline (i.e., *δ* < 0), the learned spread *S* reduces action activation, biasing choices towards safe options.

#### Theoretical analysis of prediction errors

In this section, we focus on how the participant’s reward prediction should theoretically change over the course of a trial. This is based on the theory of temporal difference learning (Sutton and Barto 2018), which has been applied to describe dopaminergic responses to rewards (Schultz, Dayan et al. 1997). The basic assumption of these theories is that participants maintain a prediction of upcoming rewards at all times. This prediction is based on learned estimates *Q* of average rewards associated with the four stimuli.

At the beginning of a trial (before the stimuli appear), participants do not have any specific information to base their prediction on. Depending on whether they anticipate the appearance of the stimuli or not, they might either predict the average reward over all trials (which we take to be the average learned value across all options), or no reward at all. Neither of these two possibilities can be ruled out on theoretical grounds only. We will therefore consider them both, and develop two versions of our hypothesis, ultimately resulting into two slightly different models (see section Results/Models).

After the options appear, participants should adjust their reward prediction based on the learned values of the displayed options. We take the participants’ updated prediction to be the average learned value over the presented options. The updated prediction should differ across conditions: if the participants learned accurate estimates of the values, their prediction would be 60 points in the both-high condition and 40 points in the both-low condition.

This change in participants’ reward expectation through the appearance of reward-predicting stimuli constitutes a prediction error called **stimulus prediction error** *δ*_*stim*_, previously described in (Jang, Nassar et al. 2019), and should cause phasic DA activity (Schultz, Dayan et al. 1997). If participants’ initial reward expectation is the average over all trials (this corresponds to the first version of our hypothesis), the magnitude of that prediction error is given by

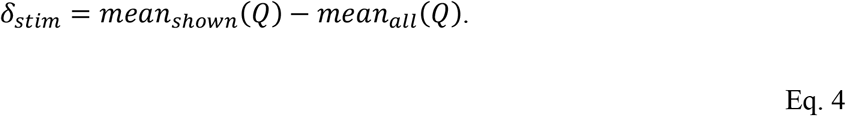

If the values of the stimuli are learned with reasonable accuracy, this prediction error should be 60 − 50 = 10 in the both-high condition and 40 − 50 = −10 in the both-low condition.

If, on the other hand, participants don’t predict any reward at the beginning of the trial (this corresponds to the second variant of our model), the magnitude would be given by

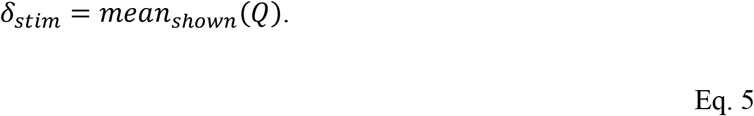

We would then expect a magnitude of 60 in the both-high condition and a magnitude of 40 in the both- both-low condition.

Next, participants will make a choice. Now, their reward expectation is the value of the option they chose. Finally, a reward is displayed, forcing participants to update their reward estimate again. This second prediction error—the difference between the learned value of the option and the actual reward received— we call the **outcome prediction error** *δ*_*out*_. Its magnitude is given by

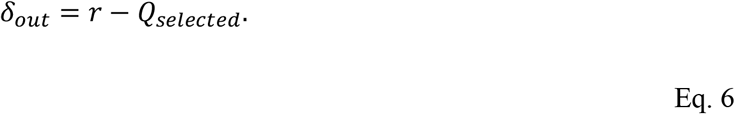

It is the outcome prediction error that drives learning about the stimuli (see Eq. 2 and Eq. 3 above).

In summary, our analysis reveals that two prediction errors (and two corresponding DA responses) should occur in each trial: first the stimulus prediction error shortly after the presentation of the options, and second the outcome prediction error after the presentation of the reward.

#### Behavioral Predictions

Here, we combine the results of the analyses above to derive task-specific behavioral predictions from our theory (see Fig 1C for a schematic representation). Let us first assume that the stimulus prediction error is given by Eq. 4. We have seen that in the matched-mean conditions (both-high and low), the presentation of the options should cause a prediction error (positive and negative, respectively), and hence a transient change in DA levels (increase and decrease, respectively) in the striatum during the choice period (Fig 1C, mechanistic level). We have also seen that DA levels affect choices through modulation of the BG pathways: increased DA makes people risk seeking, decreased DA makes them risk averse. If the average reward is similar for two options (as it is in the both-high and the both-low condition), these risk preferences should be the decisive factor in decisions (Fig 1C, task level and past rewards level).

Taken together, these premises suggest that we should see a preference for the risky stimulus in the both- high condition (depicted in Fig 1C), and a preference for the safe stimulus in the both-low condition. These effects should appear gradually, since they require that both mean and spread of the reward distributions are learned. Specifically, risk preferences in the both-high and both-low conditions should appear slower than value preferences in the difference condition. This follows from the underlying plasticity rules: one can show that the learning rate for spread must always be lower than the learning rate for value (Möller and Bogacz 2019). In addition to this, a reasonably accurate value estimate is required for the spread estimate to converge; this also contributes to a higher in learning speed for value compared to spread.

Relaxing now our assumption that the stimulus prediction error is given by Eq. 4 and allowing stimulus prediction errors of the type of Eq. 5, we can still be sure that the stimulus prediction error would be stronger in the both-high condition than in the both-low condition. We should thus see a difference in risk preference between conditions independent of the exact nature of the stimulus prediction error.

In summary, we have derived two predictions: 1) We should see a difference in risk preference between conditions, and 2) we should see gradually emerging risk seeking in the both-high and risk aversion in the both-low condition (prediction 2 depends on a stronger assumption on the nature of the stimulus prediction error). To arrive at these predictions, we took into account neural mechanisms. However, the predictions are purely on the level of behavior. In the empirical part of this study, we hence focus on behavior, conscious that this will only provide indirect evidence for the neural underpinnings of the proposed mechanism. However, it might reveal a previously unknown effect with a clear, plausible biological explanation. We provide further predictions based on neural signals in the Discussion.

### Behavior

In the previous section, we introduced a novel reinforcement learning task and performed a theoretical analysis to derive behavioral predictions. In this section, we present the results of testing these predictions.

We recruited a cohort of participants (N=27, 3 excluded, see Fig S1) and recorded their behavior in the task described above. Each participant performed four blocks of 120 trials. During each block, all six possible stimuli pairings occurred equally often. Each block used a new set of four stimuli, mapped to the same four distributions.

First, we investigated whether participants’ performance improved during the task. Based on the premise of associative learning, we expected choice accuracy (i.e., likelihood of choosing the option with the higher average reward) to increase gradually over trials. To confirm this, we focused on choices in the difference condition, where participants had to choose between stimuli with different average rewards. We found that indeed, the probability of choosing the stimulus with the higher average reward increased gradually over trials across the population. Average performance differed from chance level with high significance (Fig 2A, t-test, t(27) = 31.9, p < 0.001) and approached its asymptote in the second half of the block. These findings suggest that our participants successfully used associative learning in our task, confirming a basic assumption of our theory.

**Fig 2:**
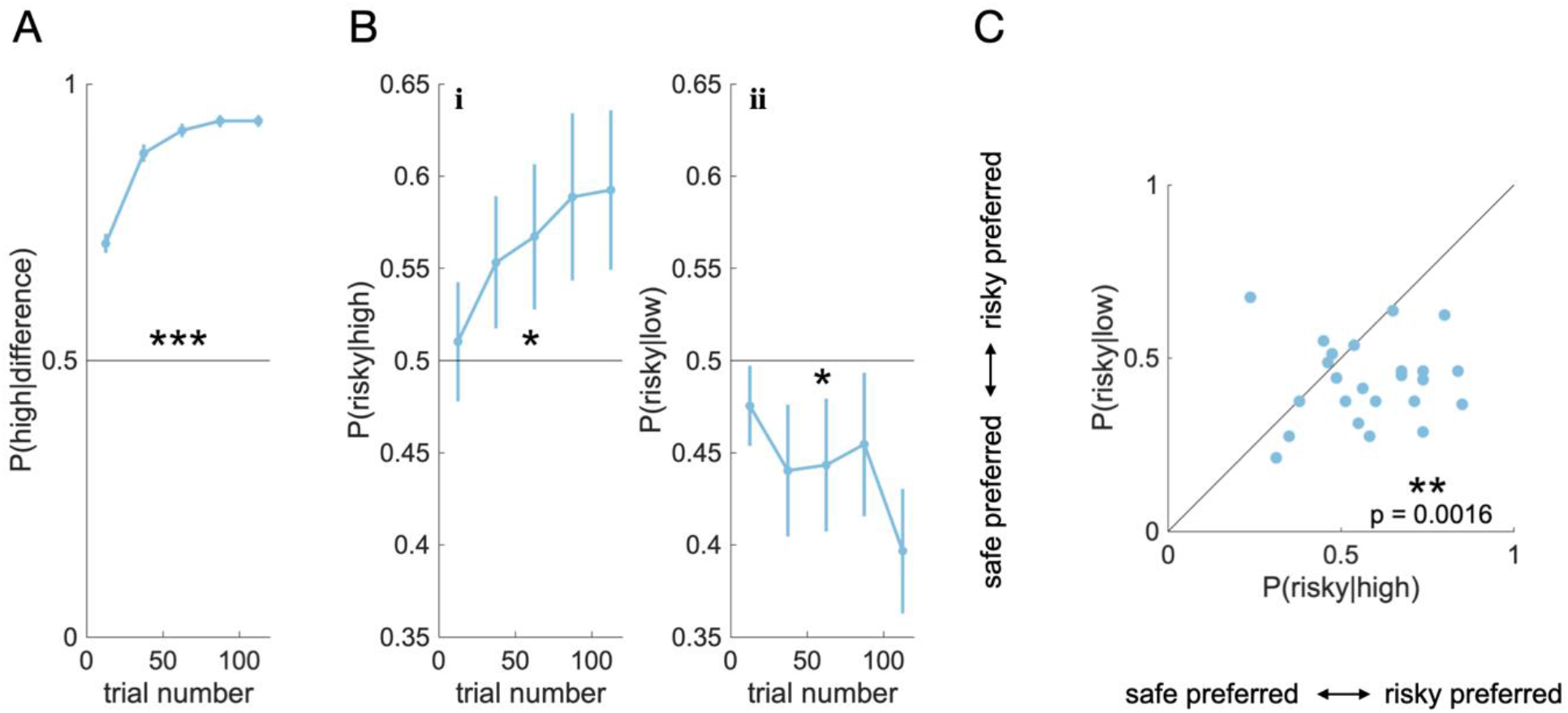
Choice probabilities A) Probability of choosing a stimulus with high average reward over a stimulus with low average reward, as a function of trial number. Choices were binned according to trial number. For each bin, we show the mean (dot) and SE (bar) over subject means. The stars (*: p < 0.05, **: p < 0.01, ***: p < 0.001) indicate that the population mean across all trials is significantly different from indifference (0.5). B) Probability of choosing the risky stimulus over the safe stimulus, i) if mean rewards are both high, ii) if mean rewards are both low, as a function of trial number. Data represented as in A. C) Correlation between risk preferences in the both-high mean and the both-low condition. Each point represents one participant. Preference for the risky stimulus if mean rewards are high is plotted against preference for the risky stimulus if mean rewards are low. If a point falls below the diagonal, the participant was more risk seeking for high-mean stimuli than for low-mean stimuli. The stars indicate that the population mean is significantly below the diagonal.

Next, we tested our first prediction—that participants would be risk-seeking in the both-high condition and risk-averse in the both-low condition, and that these effects would emerge gradually and more slowly than increases in task performance. For this, we analyzed choices in the both-high and both-low condition. For each condition, we investigated the likelihood of choosing the stimulus with the broad distribution (risky) over the stimulus with the narrow distributing (save). Preferring risky over safe was considered risk-seeking, preferring safe over risky was considered risk-averse. As predicted, we found significant risk-seeking in the both-high condition (Fig 2Bi; two-tailed t-test: p = 0.0343, t(26) = 2.23) and significant risk aversion in the both-low condition (Fig 2Bii; two-tailed t-test: p = 0.0317, t(26) = - 2.27). These preferences emerged gradually as a function of trial number, at a slower rate than the preferences for the high over low value stimuli (compare Fig 2A against Fig 2Bi and Fig 2Bii; mixed effects model (see methods): p = 0.001). In summary, the data confirmed our first prediction in all aspects.

We then tested our second prediction—that there would be a difference in risk-preferences (i.e., in the likelihood to choose the risky option over the save option) between the both-high and the both-low condition. Specifically, we predicted higher risk-seeking in both-high than in low. To test this, we computed the difference in risk-preference between conditions for each participant. We found that most of the participants were more risk seeking in the both-high condition than in the both-low condition (Fig 2C; two-tailed paired t-test: t(27) = 3.58, p = 0.0016), which confirmed our second prediction. In summary, in data recorded from the task described above, we found the two behavioral effects we predicted. The second effect was clearer than the first effect (t = 3.58 versus t = 2.23 and t = −2.27). This is consistent with our theory: the first effect rests on more assumptions than the second. We will thus focus on the second effect in the analyses below.

Our findings provide initial evidence for a behavioral link between prediction errors and risk preferences. In the next section, we use computational modelling to compare our explanation of the measured effects to alternative explanations.

### Modelling

In the previous section, we have shown that effects like those predicted by our theory can be found in experimental data. Our analysis so far rested on the assumption that participants know the ground-truth means and standard-deviations of the reward distributions. However, these statistics have to be learned when participants perform the task. To capture this learning process trial-by-trial learning models can be fit to the data. These models allow us to answer some other important questions that have not yet been addressed:

1. Are there alternative explanations for the observed effects?
2. Does our theory fit the data better than existing theories?

In this section, we use computational modelling to answer these questions. To test our theory against alternative explanations, we use simulations as well as model comparison techniques (Palminteri, Wyart et al. 2017).

#### Models

Associative learning in tasks like ours is commonly described with the Rescorla-Wagner (RW) model (Rescorla and Wagner 1972). All models we use here are variants of this base model. RW is also the first type of explanation for the effects we observed—it has been shown that even basic associative learning can yield risk preferences through sampling biases (Niv, Edlund et al. 2012).

The second type of explanation involves the utility of reward points: risk aversion as well as risk seeking have been explained as consequences of nonlinear utility functions (Kahneman and Tversky 2013). For our analysis, we considered RW-type learning in combination with the two most common families of utility functions: the concave utility of expected utility theory and s-shaped utility of prospect theory. The corresponding models are called concave UTIL and s-shaped UTIL.

The third type of explanation is based on RW with variable learning rates. It has been observed that humans use different learning rates for positive and negative outcomes (Gershman 2015), and that this can lead to risk preferences (Niv, Edlund et al. 2012). We implement this in the model pos-neg RATES. It has also been observed that when tracking a reward signal, humans adapt to the signal statistics (Daw, O’doherty et al. 2006), akin to the Kalman filter used in engineering. In Kalman filters, the learning rate depends on the variance of the signal that is tracked. We thus included a model with different learning rates for high variance and low variance rewards, referred to as variance RATES.

Finally, the explanation that we propose in this study is that prediction errors induce transient risk preferences, as detailed above. We represent our hypothesis with two very similar models, called PEIRS (Prediction Errors Induce Risk Seeking) and PIRS (Predictions Induce Risk Seeking). These models differ only in how the stimulus prediction error is computed—PEIRS assumes that the initial reward expectation corresponds to the average reward over all options (Eq. 4), while PIRS assumes that the initial reward expectation is zero (Eq. 5).

#### Alternative explanations

First, we wanted to know which of the models could reproduce (and hence explain) the observed effects. To check this, we first fitted all models to our dataset and extracted maximum-likelihood parameters for each participant. We then used these parameters to simulate behavior in our task with all candidate models to generate synthetic datasets. Finally, we analyzed these synthetic datasets in the same way as the experimental dataset. In our analysis, we focused on the difference of risk preferences between conditions (Fig 2C), as this was the most pronounced of the predicted effects.

We found that the models that represented our hypothesis reproduced the effect best: the PEIRS model captures the overall distribution of risk preferences across the population best (Fig 3A, panel PEIRS, see Fig S2 for likelihood ratios) and produces the most realistic effect size among all models we considered (Fig 3B, empirical effect size: 0.122, simulated effect size with PEIRS model: 0.087, no significant difference between experimental data and simulated data, two-tailed paired t-test, t(26)=-1.72, p=0.0976). The PIRS model, on the other hand, is the only model that captures the part of our population that is risk seeking in both the both-high and the both-low condition (Fig 3A, panel PIRS, upper right quadrant, see Fig S2 for likelihood ratios). It also produces the second most realistic effect size (Fig 3B, empirical effect size: 0.122, simulated effect size with PEIRS model: 0.059, no significant difference between experimental data and simulated data, two-tailed paired t-test, t(26)=-2.01, p=0.0552).

**Fig 3:**
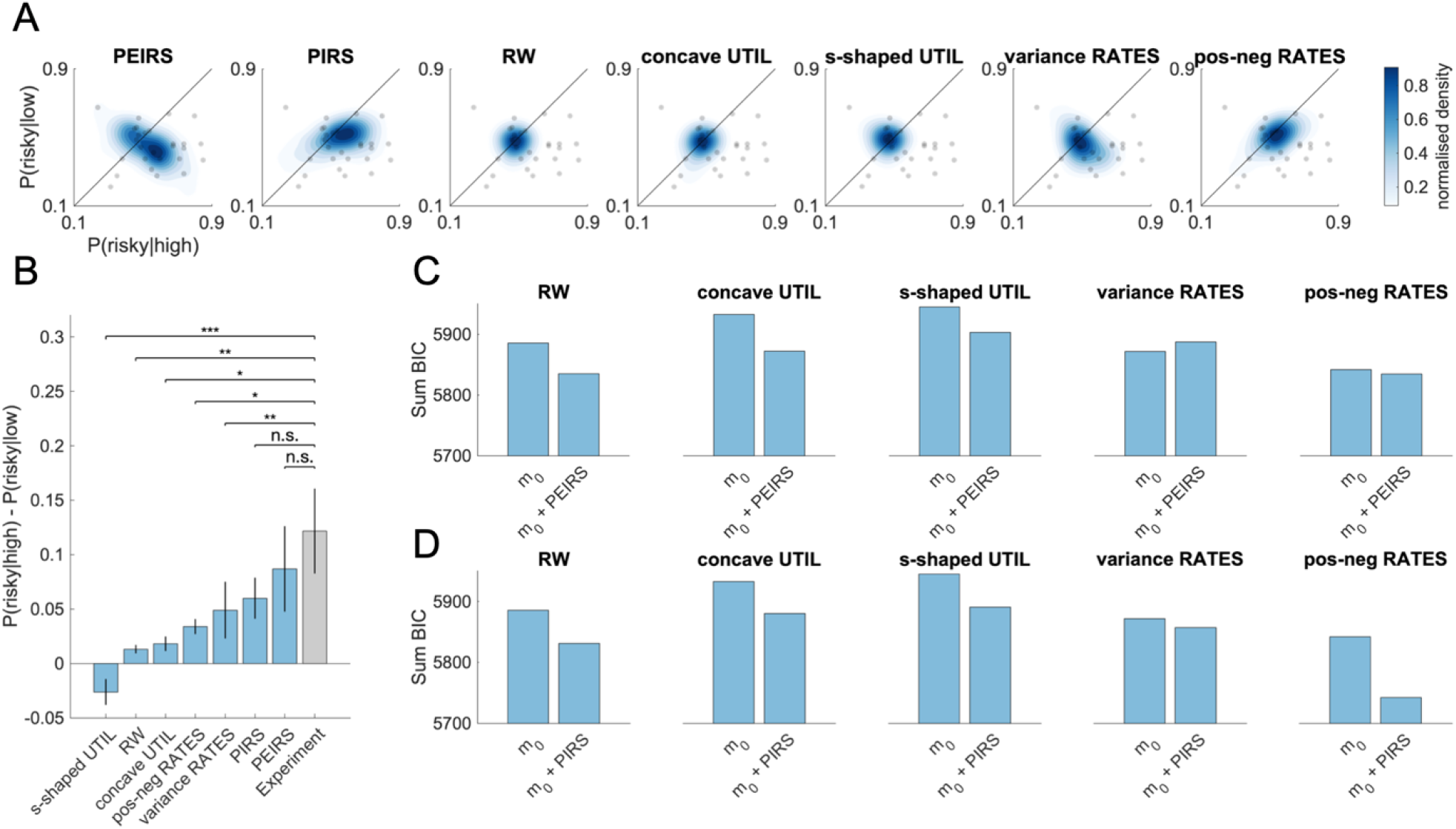
Modelling results A) Risk preferences in simulated datasets. The risk preference distributions extracted from simulated data are plotted as blue contour plots. The corresponding experimental data is superimposed as a grey scatterplot (identical in all panels). The shading corresponds to estimated probability density functions and is scaled to match each distribution’s density value range for better visibility. B) Difference in risk preferences between conditions, for simulated and empirical datasets. Simulated datasets are shown as blue bars, the experimental dataset is shown as a grey bar. We show the mean (bar) and the SE (errorbar) of subject means (averaged over experiment repetitions for simulated datasets). C, D) Model comparison with and without PEIRS (C) / PIRS (D) extension. Lower BIC indicates a better fit. The y-axis is valid for all plots in the same row.

Most of the alternative models (RW, concave UTIL, s-shaped UTIL, pos-neg RATES) predicted very narrow distributions for the risk preferences, which is at odds with our empirical results. The variance RATES model was the only alternative model that produced a somewhat realistic distribution of risk preferences (albeit not as realistic as the PEIRS model, see Fig 3A, PEIRS and variance RATES and Fig S3 for likelihood ratios).

With respect to our main effect—the difference of risk preferences between conditions—all alternative models predicted effect sizes that differed significantly from the empirical findings (Fig 3B, two-tailed paired t-tests, s-shaped UTIL: t(26)=-4.71, p < 0.001, RW: t(26)=-2.78, p=0.010, concave UTIL: t(26)=- 2.66, p=0.013, pos-neg RATES: t(26)=-2.4, p=0.024, variance RATES: t(26)=-3.04, p=0.005).

We concluded that among the tested models, our theory (specifically the PEIRS variant) was the best explanation for the observed risk preferences (both distribution and difference between conditions). None of the other models provided a convincing alternative explanation.

#### Model fit

After showing that the PEIRS model can reproduce the risk preferences that we measured, we wanted to use model comparison techniques to test more formally whether our dataset provides evidence for or against the PEIRS / PIRS models. One might be tempted to do this by simply comparing all candidate models directly and selecting the best fitting one as the winner. However, this is not the right strategy here. Consider for example a population with strongly concave utility functions and, in addition, moderate PEIRS. A direct comparison between a utility model and a PEIRS model would favor the utility model for this population, even though PEIRS was present. More generally, if it is possible that more than one of the considered effects (nonlinear utility, variable learning rates, PEIRS / PIRS) is present in the same participant, a direct model comparison between the effects will only determine the largest effect. It cannot tell us whether effects are there or not.

The correct way to test whether there is PEIRS in our dataset is to compare models with PEIRS (extended models) against models without PEIRS (base models). This should be done for different base models, as they constitute different alternative explanations. We therefore performed a fit for all conventional models (RW, concave UTIL, s-shaped UTIL, pos-neg RATES, variance RATES), and for each fit computed the Bayesian Information Criterion (BIC) across the entire population. We then added the PEIRS mechanism to each of these models, performed a fit with the extended models, and again computed the BIC across the population. Finally, we compared the BICs with and without the PEIRS extension. We repeated the same analysis with PIRS as well.

We found that adding the PEIRS extension improved parsimony for all but one model (variance RATES, Fig 3C), and that adding PIRS improved all models (Fig 3D). This suggests that PIRS and PEIRS tend to improve model fits enough to compensate for increased complexity, and hence capture some non-trivial feature of the data that the other models cannot capture.

We concluded that in all but one case, our theory helps to explain empirical data better than existing theories. The variance RATES model was not improved by PEIRS, perhaps because it already reproduced the distribution of risk-preferences fairly well (Fig 3A, variance RATES). However, variance RATES predicted an effect size that differed significantly from the empirical one (Fig 3B) and was improved by the PIRS extension (Fig 3D). This suggests that variable learning rates cannot capture all features within the realm of our theory.

Finally, we considered the possibility that prediction errors might influence choices even across trials— that the outcome prediction error of one trial might induce risk preferences in the next. To test this, we defined another model, called Outcome Errors Induce Risk Seeking (OEIRS) and performed an analysis similar to Fig 3C and 3D. We found no evidence for across trial effects, likely due to comparatively large delays (Fig S3, see supplementary discussion). This negative result is interesting in itself, but also provides additional confidence for our positive results for PEIRS and PIRS—these models did not simply win because they had the more parameters (PEIRS, PIRS and OEIRS have the same number of parameters), but because they explained the data better.

### Pupillometry

Above, we used choice data to look for the effects we predicted, and computational modelling to rule out alternative explanations. In this section, we use an additional physiological marker—pupil dilation—to provide additional evidence for our hypothesis. It is known that pupils dilate in response to surprise (Preuschoff, t Hart et al. 2011, Cavanagh, Wiecki et al. 2014, Browning, Behrens et al. 2015, Lawson, Mathys et al. 2017). This means that they allow an indirect readout of the magnitude of prediction errors, since outcomes that are further from the prediction are more surprising.

We exploit this in three steps: first, we confirm that we can read out the magnitude of prediction errors by computing the pupil response to the well documented outcome prediction errors. Second, we prove the occurrence of stimulus prediction errors by extracting the corresponding pupil response. Finally, we show that the strength of the stimulus prediction error, as measured through pupil dilation, is connected to the strength of risk seeking extracted from choices.

#### Pupil dilation reflects outcome prediction errors

Can we measure prediction error magnitude through pupil dilation? To test this, we took the absolute value of the outcome prediction error |*δ*_*outcome*_| as a measure of surprise. Trial-by-trial estimates of this prediction error were extracted from the PEIRS model fits. Regression analyses were used to determine whether pupil dilation after reward presentation encoded |*δ*_*outcome*_|. In this analysis, we focused on the first half of the block (trials 1 to 60) where the learning from feedback occurs (Fig 2A). We found a phasic response which peaked 0.9 s after reward presentation (Fig 4A t-test: t(26) = 4.61, p = 0.0001; statistical significance was established through leave-one-out unbiased peak detection and confirmed through a cluster-based permutation test, see Fig S4A). This confirmed pupil dilation as a proxy for prediction error magnitude.

**Fig 4:**
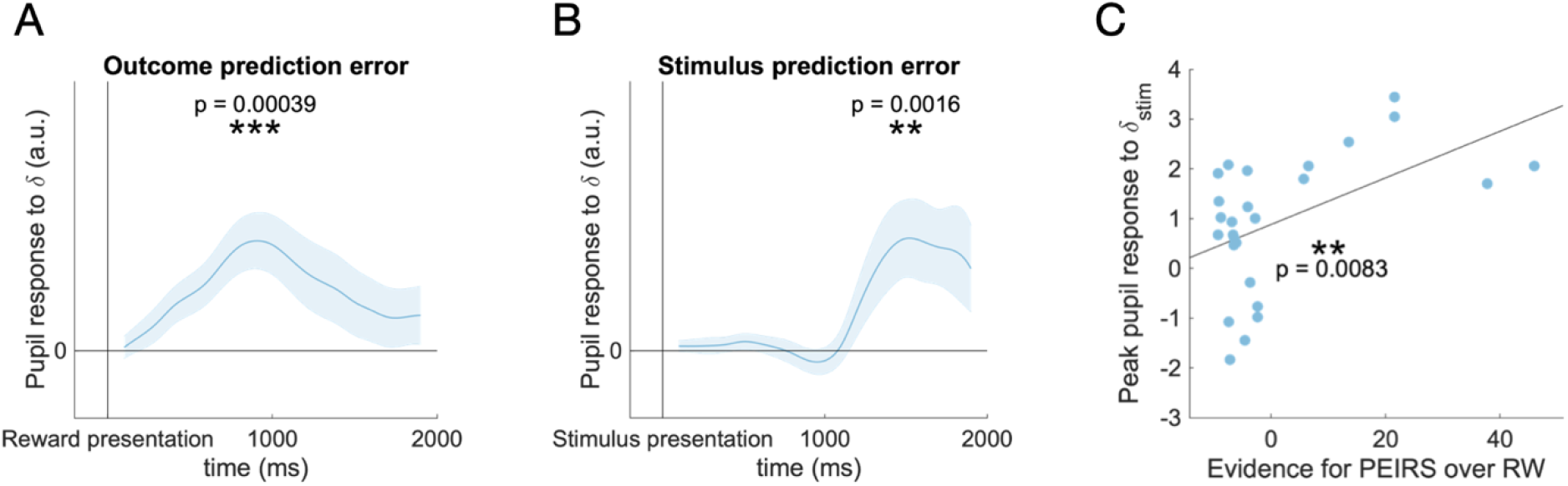
Pupil responses to prediction error magnitude index behavioral effects A) Pupil dilation encodes the magnitude of the outcome prediction errors in the first half of the block (trials 1 −60). The blue line and shading represent mean and SE across participants of the regression coefficients obtained from a linear model. Pupil dilation was predicted from trial-by-trial estimates of outcome prediction errors, obtained from the fitted PEIRS model. Responses were aligned at reward presentation. For display, the trace was smoothed using spline interpolation. B) Pupil dilation encodes the magnitude of the stimulus prediction errors in the second half of the block (trials 61 - 120). Similar to A), with pupil dilation predicted from trial-by-trial estimates of stimulus prediction errors. Responses were aligned at stimulus presentation but truncated before the time of the outcome. C) Effect strength in behavior predicts effect strength in pupil dilation. The plot shows the correlation between BIC_RW_ – BIC_PEIRS_ (x-axis) and the regression coefficient for the stimulus prediction error as a predictor of pupil dilation, at time of maximum effect strength (y-axis), which was determined using a leave-one-out procedure.

#### Pupil dilation confirms occurrence of stimulus prediction errors

Next, we sought to confirm the occurrence of stimulus prediction errors, which are essential for our theory of prediction-error induced risk seeking. For this, we extracted estimates of *δ*_*Stimulus*_ V for every trial and used an analysis similar to the previous step to extract the corresponding pupil response. Here, we aligned the pupil signal at stimulus presentation, and censored all data points collected after reward presentation to avoid confounding factors such as reward or outcome prediction errors. We also focused on the second half of the block (trials 61 to 120) where we observe pronounced risk preferences (Fig 2B). We found a phasic response which peaked 1.6 s after stimulus onset (Stimulus prediction error: Fig 4B; t-test: t(26) = 2.89, p = 0.0079. Statistical significance was established through leave-one-out unbiased peak detection and confirmed through a cluster-based permutation test, see Fig S4B). This confirms the existence of a stimulus prediction error that occurs after stimulus onset, as hypothesized.

The response to the stimulus prediction error was roughly similar to the response to the outcome prediction error, except for a longer delay between stimulus prediction error onset and the peak of the pupil response. There might be many reasons for this difference in delay. Among those, differences in information processing might play a role: generating a stimulus prediction error involves two stimuli, hence attention mechanisms, in addition to retrieval of value estimates from memory. Generating the outcome prediction error, on the other hand, just requires the processing of a number.

#### Stimulus prediction errors are linked to risk seeking

Our theory suggests that risk seeking is caused by transiently increased dopamine levels that reflect reward prediction errors, specifically the stimulus prediction error. If this is the case, we should expect to see stronger risk preferences in individuals that respond stronger to stimulus prediction errors.

We tested this using individual behavior and pupil responses. To quantify the strength of prediction error induced risk preferences in behavior, we used the difference in BIC computed from the RW fit and the PEIRS fit (see Results/Modelling). This difference is a measure of evidence for the PEIRS model over the RW model, and hence reflects the overall strength of the effects of interest. The strength of the pupil response was quantified through the regression factor’s t-statistic at the time of strongest effect (determined though leave-one-out peak detection to avoid bias). Using a linear model to relate pupil effect strength and behavioral effect strength, we found that participants responded stronger to stimulus prediction errors if they showed more pronounced prediction-error induced risk seeking in their behavior (Fig 4C, adjusted R^2^: 0.23, p = 0.008). This confirmed our prediction, thus providing additional evidence to our explanation.

## Discussion

Different behavioral phenomena—learning from prediction errors and biased risk-preferences–are attributed to the same neurotransmitter, dopamine. Based on this, we first developed a model that predicts shifts in behavioral risk preference as a function of reward prediction errors. Second, we tested it using a task where reward prediction errors are immediately followed by decisions that involve risk. We found that reward prediction errors and the probability of subsequent risk-taking are positively correlated: positive reward prediction errors induce risk seeking, negative ones inhibit it. Finally, we showed that the magnitude of the reward prediction error (as indexed by pupil dilation) determines its effect on risk-preferences.

Our results are consistent with our initial hypothesis: the two roles of dopamine (teaching signal and risk modulator) interfere with each other. This study hence provide evidence *against* the conjecture that the roles of dopamine are well separated. That conclusion fits in well with other recent findings: it has been shown that phasic dopamine correlates with motivational variables (Hamid, Pettibone et al. 2016) and movement vigor (da Silva, Tecuapetla et al. 2018) just as well as with reward prediction errors. Together, these findings cast doubt on the separation into tonic and phasic and on separations in general.

### Predictions or Prediction errors?

In our theoretic analysis, we considered two possibilities for how the stimulus PE might be calculated: the first possibility was to take the difference between value of the options and the overall average reward.

This was codified in the PEIRS model. The other possibility was to take the difference between the value of the options and zero, which corresponded to no reward expectation during the ITI. This was codified in the PIRS model, which is short for Predictions Induce Risk Seeking. We called it as such because in this case, there is no way to distinguish between prediction and prediction error. Our task design in general is not suitable to empirically differentiate between these two possibilities, because predictions and prediction errors are strongly correlated at stimulus presentation. We did not aim to empirically distinguish the two mechanisms, but rather, to show that either of these mechanisms explains our data better than alternative explanations (with one exception, see Fig 3C and D). Differentiating between the two mechanisms is an interesting direction for future work.

Another question that might arise in this context is about the role of the outcome prediction error of the previous trial. According to our theory, that prediction error should be broadcasted by the dopamine system just like the stimulus prediction error and might therefore also affect risk preferences. Of course, there is a difference in timing: the choice follows the outcome prediction error of the previous trial with 3.47 s delay on average (standard deviation 0.51). In contrast, there is only an average delay of 0.97 s (standard deviation 0.51) between stimulus onset and choice. We might thus expect that the impact of the outcome prediction error, if at all observable, might be much weaker than that of the stimulus prediction error. A supplementary analysis similar to those displayed in Fig 3C and 3D confirmed this: there is no evidence for an association of risk preferences and outcome prediction error of the previous trial in our dataset (Fig S3).

### Relation to behavioral economics

Decision making under uncertainty has been extensively studied in behavioral economics. One main finding in this field, codified in prospect theory, is that humans tend to be risk-averse if decisions concern gains, and risk-seeking if decisions concern losses (Kahneman and Tversky 2013). However, those classic findings rely on explicit knowledge about the probabilities involved in the decisions. Several more recent studies indicate that those tendencies reverse when risks and probabilities are learned from experience (i.e., by trial and error): if learning is incremental and based on feedback, humans tend to make risky decisions about gains and risk-averse decisions about losses (Wulff, Mergenthaler-Canseco et al. 2018).

This phenomenon has been termed the description-experience gap and is considered a “major challenge” for neuroeconomics (Garcia, Cerrotti et al. 2021). In cognitive neuroscience and psychology, some studies have reproduced this phenomenon (Madan, Ludvig et al. 2014), while others report risk aversion in the gain domain (Niv, Edlund et al. 2012).

Our task differs somewhat from the tasks studied in the description-experience gap literature, since we only use gains (reward points are always positive). However, it is also known that humans often evaluate outcomes with respect to a reference point (Kahneman and Tversky 2013). In the context of our task, it seems plausible that rewards are evaluated relative to the average of previous rewards, or relative to the middle of the experienced reward range, which rapidly converges to 50 points within the first few trials. Rewards under 50 (and hence the majority of the rewards that result from choosing stimuli 3 and 4) would then be considered losses. Decisions between the two low-valued stimuli (both-low condition), would fall in the loss domain. Under this perspective, our results are in line with the description-experience-gap, and differ to those of (Niv, Edlund et al. 2012). This might be due to the degree of implicitness of the knowledge that is gained during the task: Niv et al. used classical bimodal reward distributions (e.g., 40 points with probability 50 %, 0 points otherwise) which participants might be able to recognize as such after a few trials. Here, we used high-entropy reward distributions (normal distributions, see Fig. 1B), which could not be mapped onto bimodal gambles, and thus made anything but implicit learning intractable.

### Relation to other theories

For our behavioral results, interpretations other than our dopaminergic explanation may be evoked: the behavior in a similar task (Madan, Ludvig et al. 2014) was interpreted as the result of memory replay: experiences (“Obtained reward X after choosing option Y”) might not only be used for immediate value updates but might also be stored in a memory buffer. This buffer can then be used for offline learning from past experiences in times of inactivity, such as during the inter-trial interval. It was proposed by (Madan, Ludvig et al. 2014) that experiences are more likely to enter the buffer if they are extreme. If entering the buffer is biased in this way, then so are the values learned from replaying those experiences. In our task, extreme might mean that the reward was extremely high or low. The corresponding bias would drive choice towards the stimuli that produce the highest rewards, and away from those that produce the lowest, and thereby lead to a pattern similar to the one we observed.

Which theory is closer to the truth? One relevant piece of data is the gradual emergence of the risk preferences (Fig 2B). We detailed above that our theory predicts slowly emerging risk preferences. In particular, risk preferences should emerge after value preferences. The memory theory, on the other hand, holds that risk preferences stem from biased values. If that was the case, risk preferences should emerge at the same rate as value preferences, which is not what we observed.

Beyond this, it is difficult to compare the memory theory directly to prediction-error induced risk-seeking; it is unclear how to obtain trial-by-trial choice predictions from the memory model, which rules out a formal model comparison. Indeed, the memory model has so far only been fitted to and assessed based on summary statistics of a large collection of trials. Further, the memory model has so far not been equipped with a mechanistic underpinning and was therefore not validated on physiological variables such as pupil dilation. In contrast, prediction-error induced risk-seeking can be fitted trial-by-trial, allowing it to make predictions not only about summary statistics but about the evolution of preferences during the task as well as about the immediate impact of extreme events such as large prediction errors. The corresponding latent variables can be correlated with physiological variables, proving that they can explain aspects of pupil dilation in addition to behavior (Fig. 4C and 4D).

### Further experimental predictions

In this paper, we present a theory based on neural effects, and test some of its behavioral predictions. The results could have falsified our theory, but they did not—all our predictions where confirmed. This suggests that our theory is valid on the level of behavior, but it does not prove that our theory is correct on the level of neural mechanisms. More direct measurements are needed to establish this.

Correlational studies are possible. They should have sufficient time resolution to differentiate the prediction errors at different times during the trial, such as EEG, electrophysiology, voltammetry or similar. With those, one could gain more direct measurements of the PEs, and hence resolve more clearly how they are associated with subsequent risk seeking. Another possibility are causal studies. For example, optogenetic tools could be used to elicit or suppress prediction errors just before choices involving risk, for instance by stimulation VTA dopamine neurons at the time of stimulus onset.

### Conclusion

In summary, we demonstrate that a biologically inspired theory of basal ganglia learning predicts an interaction between prediction errors and risk seeking. This is based on dopamine’s dual role in learning and action selection. We present behavioral data that matches these predictions. We further show that between-participant variability in behavior can be linked to differences in pupil responses—the stronger the pupil response to stimulus prediction errors, the stronger the prediction error induced risk seeking.

## Methods

### Participants

We tested 30 participants (15 female, median age: 26, range: 18-42). Our participants did not suffer from visual, motor or cognitive impairments. Participants were recruited from the Oxford Participant Recruitment system and gave informed written consent. All experimental procedures were approved by the Oxford Central University Research Ethics Committee, approval number R45265/RE004.

Participants were given a written set of instructions, as well as an oral instruction. They were first provided with a description of the task sequence. Then, they were told that the rewards were random, but nevertheless higher on average for some shapes than for others. Finally, they were advised to get as much total reward as possible and that their compensation would be between 8 and 12 GBP, depending on their performance. After the task, all participants received a compensation of 10.50 GBP.

Our results are based on 27 of the 30 participants. Three participants were excluded from the analysis due to their failure to understand the task: we evaluated the participants’ understanding of the task by scoring their preferences in different-mean choices during the second half of the blocks. Participants were included in the analysis if they chose the high-valued option in more than 70 % of those trials (Fig S1).

### Emerging preferences

To test whether value preferences emerged faster than risk preferences, we used a linear mixed effects model to model choices. As predictors, we included fixed effects of decision type, trial number, and their interaction. Here, decision type was defined as value decision (difference condition) versus risk decision (both-high and both-low conditions). We also included random effects for all predictors and a random intercept, by participant. A likelihood ratio test for the fixed effect of the interaction between trial number and decision type revealed a significant positive effect (p < 0.001; based on comparing the empirical likelihood to a distribution of likelihoods obtained from 1000 Monte Carlo simulations of data from the model without the fixed effect, as implemented in MATLAB’s ‘compare’ function). This confirms value preferences emerged faster than value preferences (Fig 2A and 2B).

## Models

The models we used in this study are all variants of the RW model (Rescorla and Wagner 1972). They consist of learning rules for latent variables, such as value and spread, and choice rules that convert latent variables into choice probabilities. Below, we provide these rules for all models, along with some auxiliary rules as necessary.

For all models, *i* ∈ {1,2,3,4} is the stimulus index, and *Q*_*i*_ is the value of stimulus i. The index *j* is used for the options on screen in each trial (for example, *j* = {1,3} if stimuli 1 and 3 are shown). The initial values at the beginning of each block are denoted by *Q*_0_. The update rules are only applied to the values and spreads of the chosen stimuli; the values and spreads of unchosen stimuli do not change. The probability of choosing stimulus *i* is denoted by *p*_*i*_, the received reward is denoted by *r*.

### RW

In the RW model, learning is driven by differences *δ*_*outcome*_ between the reward expected and received:

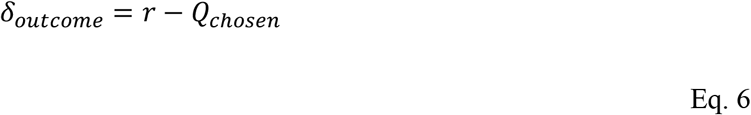

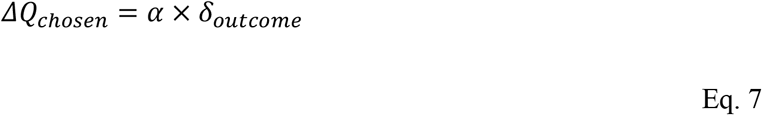

Here, *α* denotes the learning rate. Learned values were linked to choice probabilities via a standard softmax rule (Daw 2011):

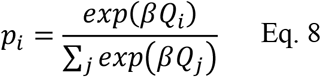

The RW model has free parameters *α* ∈ [0,1], *β* > 0 and fixed parameters *Q*_0_ = 50.

### Concave utility

To allow for concave subjective utility of reward points in our experiment, we used an exponential family of functions (Guyaguler and Horne 2001), which we adapted to our reward range through appropriate scaling (*σ*) and shifting (*m*):

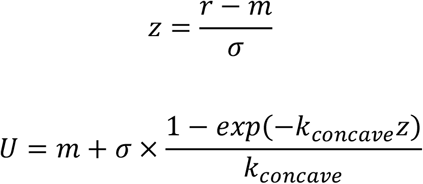

The nonlinear utility of reward enters through the computation of the prediction error:

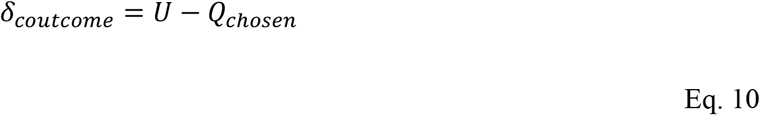

Updates are computed with Eq. 7, choices are modeled using the softmax rule in Eq. 8. The concave UTIL model has free parameters *α* ∈ [0,1], *β* > 0, *k*_*concave*_ > 0. and fixed parameters *Q*_0_ = 50, *σ* = 50, *m* = 50.

### S-shaped Util

Utilities can also be s-shaped, with different curvatures on both sides of a reference point (Kahneman and Tversky 2013). We modelled s-shaped utility functions using sign-preserving power functions (Spitzer, Waschke et al. 2017), which we adapted to our reward range:

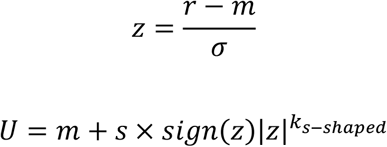

Prediction errors are calculated as in Eq. 10, updates happen according to Eq. 7, choices are modeled using the softmax rule in Eq. 8. The s-shaped UTIL model has free parameters *α* ∈ [0,1], *β* > 0, k_*s*−*shaped*_ ∈ [0,1] and fixed parameters *Q*_0_ = 50, *σ* = 50, *m* = 50.

### Different learning rates for positive and negative prediction errors

It has been shown that learning from positive outcomes can differ from learning from negative outcomes (Gershman 2015). We model this by letting the learning rate depend on the sign of the prediction error (computed according to Eq. 6). The update rules then become:

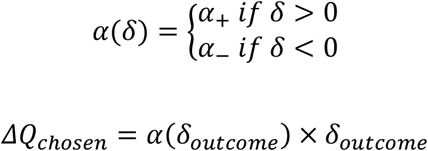

Choices are modeled using the softmax rule Eq. 8. The pos-neg RATES model has free parameters *α*_+_ ∈ [0,1], *α*_−_ ∈ [0,1], *β* > 0, and fixed parameters *Q*_0_ = 50.

### Different learning rates for noisy and save stimuli

Similarly, it has been shown that the statistics of the reward distribution can have an effect on the learning rate (Daw, O’doherty et al. 2006, Behrens, Woolrich et al. 2007). In our task, normative theory (Welch and Bishop 1995) suggests that the learning rate should depend on the variance of the signal that is being learned. We model this by allowing different learning rates for different levels of noise. The update equations become:

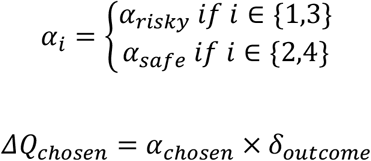

Prediction errors are computed according to Eq. 6, choices are modeled using the softmax rule in Eq. 8. The pos-neg RATES model has free parameters *α*_*ris*k*y*_ ∈ [0,1], *α*_*safe*_ ∈ [0,1], *β* > 0, and fixed parameters *Q*_0_ = 50.

### PEIRS

Our model of prediction error induced risk seeking is based on recent models of the basal ganglia (Mikhael and Bogacz 2016, Möller and Bogacz 2019). The relevant equations have been derived in section Results/Theory. They describe how the average and the spread *S*of a reward signal can be tracked. Here, *S*_*i*_ denotes the spread of the spread of the rewards received for choosing stimulus *i*. The update equations for value are given as:

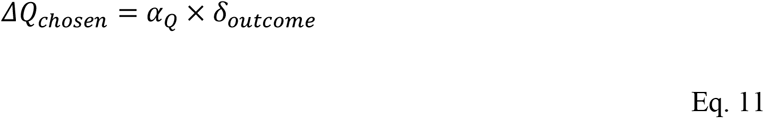

With the outcome prediction error from Eq. 6. The update equation for spread is given as:

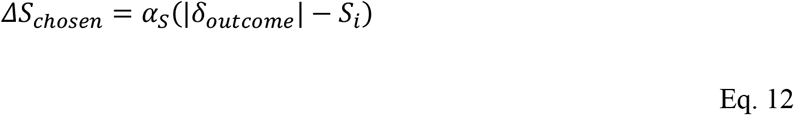

The choice rule is a variant of the softmax rule in Eq. 8:

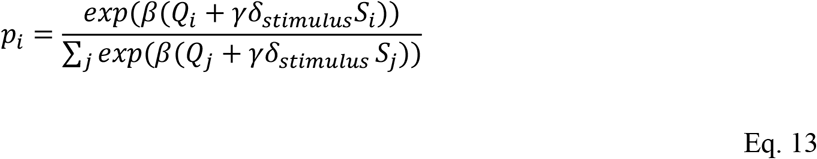

Here, spreads can contribute to decisions. The impact of spread on choice probability is gated by the stimulus prediction error:

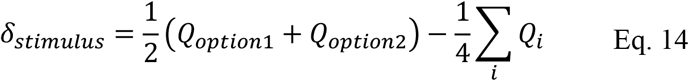

The PEIRS model has free parameters *α*_*Q*_ ∈ [0,1], *α*_*S*_ ∈ [0,1], *β* > 0, *γ S*_0_ > 0 and fixed parameters *Q*_0_ = 50. The initial spread estimate *S*_0_ was allowed to vary since participants were not given no prior information about the spread magnitude. Initial estimates might thus have differed strongly across individuals.

### PIRS

In another variant of the PEIRS model predictions induce risk seeking. It differs from PEIRS only in how the stimulus prediction error is computed:

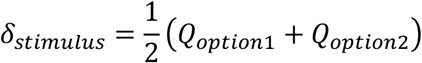

The above formula is used instead of Eq. 14. Updates and choices are governed by Eq 11, 12 and 13. The PIRS model has free parameters *α*_*Q*_ ∈ [0,1], *α*_*S*_ ∈ [0,1], *β* > 0, *γ, S*_0_ > 0 and fixed parameters *Q*_0_ = 50.

### OEIRS

If risk-preferences are associated with stimulus prediction errors, they might also be associated with outcome prediction errors from the previous trial. This is the assumption of the OEIRS model, which differs from the PEIRS model in that the outcome prediction error *δ*_*outcome*_ (computed according to Eq. 6) is substituted for *δ*_*stimulus*_ in Eq. 13. Otherwise, OEIRS is identical to PEIRS and PIRS.

### Parameter transformations and Priors

We used exponential and sigmoid transformations to constrain the parameters to their appropriate ranges. Priors were specified as multivariate normal distributions over the untransformed parameters. All but diagonal elements of the covariance matrices of those normal distributions were set to zero. Hence, the prior distributions could be factorized into univariate normal distributions (one for each parameter). Below, we provide the statistics of those prior distributions. For parameters that occur in more than one model (such as learning rate *α*), we used the same priors across models. The notation *X*∼*N*(*μ*, *σ*) means that *X* has a normal distribution with mean *μ* and variance *σ*.

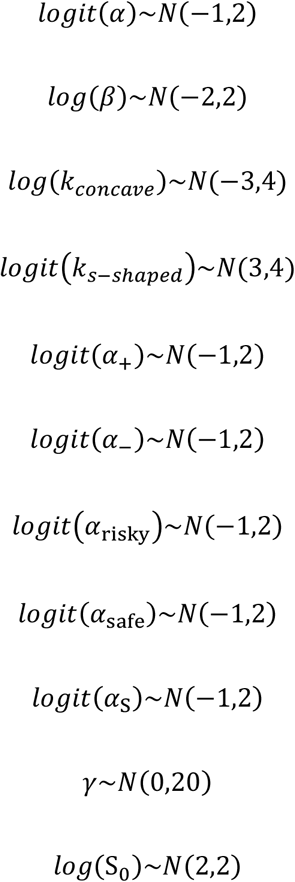

#### Fitting & Simulation

Fits were performed using the VBA toolbox (Daunizeau, Adam et al. 2014). This toolbox implements a Variational Bayes scheme. It takes a set of measurements, a generative probabilistic model that describes how the measurements arise (which usually contains some latent, i.e., unobserved, variables) and prior distributions over the model parameters as input, and outputs among other things an approximate posterior distribution over model parameters, an approximate posterior distribution over the latent variables, and fit statistics such as the BIC. We estimated parameters and latent variables (such as values and prediction errors) using the mean of the posterior distributions over parameters from the toolbox outputs.

Simulations were performed with parameters taken from fits: for each model, a fit provided us with 27 parameter sets corresponding to the 27 participants. Using those parameter sets we simulated our task and generated datasets of the same size as the dataset we obtained in our experiment. We repeated this 1000 times to obtain stable distributions. We extracted 27000 pairs of risk preferences per model from the simulated datasets (27 simulated participants in each of 1000 simulated experiments). We then used a kernel smoothing function to estimate the probability density underlying the distribution of risk preferences (MATLAB’s *ksdensity* function). Finally, we computed and visualized isolines of those estimated probability densities using MATLAB’s *contour* function (Fig 3A).

For Fig 3B, for each model we first computed the average difference in risk preference between conditions for each simulated participant in each simulated experiment, yielding 27000 differences in risk preference per model. We then averaged those across experiments, obtaining 27 differences in risk preferences per model. Those distributions were compared to the empirical distribution of mean difference in risk preference between conditions, which has 27 datapoints as well (one for each participant).

### Pupillometry

We recorded time series of pupil diameters for every trial, using an EyeLink 1000 system. For each participant, the system was calibrated before the first block, and after subsequent blocks if required.

The raw measurements were screened for blinks, which were identified as missing values in the recordings. Data points in the direct vicinity of blinks as well as data segments shorter than 50 ms were removed. After cleaning, missing value segments of less than 400 ms were filled using linear interpolation (Manohar 2019). Then, the traces were aligned to the relevant temporal markers (stimulus onset, or reward onset). We used the mean over 500 ms prior to the alignment point to define a trial-wise baseline. All traces were divided and shifted by that baseline, resulting in traces reflecting the relative change of pupil diameter after the alignment point. Finally, traces were downsampled to 10 Hz.

Pupil responses to prediction errors were obtained using linear models: for each participant, we regressed pupil dilation against a trial-by-trail estimate of the prediction error magnitude of interest, which was obtained from a model fit. This was done for each time bin of the pupil signal and resulted in a pupil response time course (beta weights and t-statistics as functions of time) for each individual. The population response was computed as the average of individual responses.

To uncover the pupil response to the stimulus prediction error, we aligned the pupil time courses at stimulus onset. After stimulus onset, participants would eventually make a choice (with variable delay, the median reaction time was 0.86 s) and receive a reward (with a 1 s delay) after their choice. Since the reward or the resulting outcome prediction error might confound our regression analysis, we censored out all data after reward presentation. This means that the number of observations on which regressions can be based rapidly declines after the median reward presentation time, which is at 1.86 s after stimulus onset. Estimates obtained later are increasingly unreliable, since they are based on insufficient data. We hence conducted our analyses for the interval 0 s to 1.9 s after stimulus onset. This allows us to obtain reliable estimates of the statistics, while still avoiding confounding effects related to reward presentation.

To test whether the pupil responses to the prediction errors are statistically significant, we performed a test at a single time-point corresponding to the peak of the effect. To avoid circularity, the time of the peak was identified using a leave-one-out method: For each participant, we used the individual pupil response time courses of all other participants to determine the time of the peak. This was achieved by executing t-tests on the response strength (t-statistic) across participants in each time bin and selecting the bin with the smallest p-value. We then took the left-out participant’s response strength from that time bin, considering it to be their response strength at the peak of the group response. In a final step, we pooled all those individual response strengths and used a t-test to check whether they deviated significantly from zero.

## Notes

### Competing Interest Statement

The authors have declared no competing interest.

